# The genome sequence of *Hirschfeldia incana*, a species with high photosynthetic light-use efficiency

**DOI:** 10.1101/2022.01.29.478283

**Authors:** Francesco Garassino, Raúl Y. Wijfjes, René Boesten, Frank F. M. Becker, Vittoria Clapero, Iris van den Hatert, Rens Holmer, M. Eric Schranz, Jeremy Harbinson, Dick de Ridder, Sandra Smit, Mark G. M. Aarts

## Abstract

Photosynthesis is a biophysical and biochemical process that plays a key role in sustaining plant and human life, being the first step in the production of energy-rich molecules and oxygen in the biosphere. Improving the photosynthetic capacity of agricultural crops is highly desirable to increase their yields. While the core mechanisms of photosynthesis are highly conserved, certainly in higher plants, plants that can maintain a high photosynthetic light-use efficiency at high irradiance are exceptional and may be useful to understand and improve high irradiance photosynthesis of crops. One such exceptional species is *Hirschfeldia incana*, a member of the well-studied Brassicaceae family that is easy to grow under standard laboratory conditions, providing an excellent resource for studying the genetic and physiological basis of this trait. Here, we present a reference assembly of *H. incana* and affirm its high photosynthetic efficiency relative to the Brassicaceae species *Brassica rapa, Brassica nigra*, and *Arabidopsis thaliana*. We estimate that it diverged from *B. rapa* and *B. nigra* 10-11 million years ago and that its genome has diversified from that of the latter two species through large chromosomal rearrangements, species-specific transposon activity, and differential retention of duplicated genes. Genes present at copy numbers different from *B. rapa* and *B. nigra* include those involved in photosynthesis and/or abiotic stress, which may mediate the high photosynthetic efficiency of *H. incana*. We expect the reference assembly of *H. incana* to be a valuable genomic resource for identifying ways to enhance photosynthetic rates in crop species.

## Introduction

Photosynthesis is the biophysical and biochemical process that sustains most life on planet Earth. The most common form of photosynthesis, oxygenic photosynthesis, uses solar energy to convert the inorganic carbon dioxide (CO_2_) to organic carbon, typically represented as a carbohydrate, releasing molecular oxygen (O_2_) from water in the process. Terrestrial plants provide by far the most conspicuous example of oxygenic photosynthesis (referred to as photosynthesis from now on for brevity) and are responsible for about 50% of the primary production of oxygen in the biosphere, with marine production by eukaryotic algae and cyanobacteria comprising the other 50%. Agriculture depends on primary production by plants, so expanding our knowledge of photosynthesis is crucial if we are to meet many of the pressing global challenges faced by mankind.

One of these challenges is the need to substantially increase the yield of agricultural crops to meet the increasing demand not only for food and fodder, but also for fibers and similar plant products, and organic precursors for the chemical industry as it transitions away from fossil carbon sources. A major yield-related trait is the conversion efficiency of absorbed solar irradiance to biomass (*ϵ_c_* [1]), a parameter which is strongly influenced by the light-use efficiency of photosynthesis. As light intensity, or irradiance, increases, the photosynthetic light-use efficiency of leaves and other photosynthetic organs decreases, which leads ultimately to the light-saturation of photosynthesis [2, 3, 4, 5, 6, 7]. Once light-saturation is reached, any additional light will not lead to any further increase in the photosynthetic rate and may even be detrimental to photosynthesis. For most crops, this light-saturation phenomenon is one of the few aspects of photosynthesis which has not yet been optimized in order to increase yield and the threshold for light saturation generally lies far below the maximum level of irradiance experienced in the field or in the greenhouse [8]. Improving the photosynthetic light-use efficiency of crop plants thus paves the way towards increasing their *ϵ_c_* and ultimately their yield [9, 8, 10, 11, 12].

The means with which to reduce the loss of photosynthetic light-use efficiency in crop plants might already exist in nature. Most crop species make use of the C_3_ photosynthetic pathway, which is the original and ancestral pathway in higher plants, with the alternative CAM and C_4_ pathways having evolved as an adaptation to heat and drought, and low CO_2_ levels and drought, respectively. Due to several issues associated with the C_3_ pathway, the maximum photosynthesis rates commonly observed among C_3_ species are generally lower than those of C_4_ ones. Although the core mechanisms of photosynthesis are highly conserved [13, 14], natural variation in photosynthesis rates has been observed for major crops such as wheat [15], rice [16, 17], maize [18], soybean [19], sorghum [20], as well as for the model species *Arabidopsis thaliana* [21, 22]. Much higher levels of variation can be expected in other species that are more ecologically specialized [23]. For example, species growing in the Sonoran Desert such as *Amaranthus palmeri, Chylismia claviformis, Eremalche rotundifolia*, and *Palafoxia linearis* were observed to sustain exceptionally high light-use efficiencies (and high assimilation rates) at high irradiance [24, 25]. Although data collected on these species provided clues about the anatomical and physiological basis of their high photosynthesis rates [25, 26], a comprehensive ecophysiological explanation of their phenotypes is still missing.

To understand the physiological and genetic basis of this more efficient photosynthesis at high irradiance, a suitable model species is needed. To date, of the handful of species showing high light-use efficiency that have been described [24, 25], none would qualify as a model species due to a combination of complex genetics and difficulties in growing in laboratory conditions (*e.g*. difficult seed germination). Taking inspiration from *A. thaliana*, an attractive model species for high light-use efficiency would need to be easily grown in regular and high-light laboratory conditions, have a high-quality reference genome, be a diploid species capable of producing a large number of progeny (hundreds of seeds from a single mother plant) with a short generation time, and allow for both inbreeding and outcrossing [27, 28].

*Hirschfeldia incana* (L.) Lagr.-Foss. is an excellent candidate that fulfils these requirements. *H. incana* is a thermophilous and nitrophilous annual species native to the Mediterranean basin and the Middle-East, but currently widespread in most warm-temperate regions of the world [29]. It is generally self-incompatible and thus allogamous, but a degree of self-compatibility has been observed in natural populations [30]. Although it makes use of the C_3_ pathway, *H. incana* has a very high photosynthesis rate at high irradiance [31], much higher than that of the C_3_ crop species wheat [15] and rice [17], more in the range of C_4_ species [32, 33]. Besides its exceptional physiological properties, *H. incana* is also an attractive model species for practical and genetic reasons. It shows fast and sustained growth in laboratory conditions and is a member of the Brassiceae tribe within the well-studied Brassicaceae family, allowing the transfer of many genetic and genomic resources developed for the model species *A. thaliana* and its close relatives *Brassica rapa* [34, 35, 36, 37, 38], *Brassica nigra* [39, 40], *Brassica oleracea* [41, 42, 37], and *Brassica napus* [43, 44]. Yet, *H. incana* has received little attention from the research community so far, being recognised mainly as a possible lead (Pb) hyperaccumulator [45, 46, 47] and for the ecological implications of its occurrence as a weed [48, 30, 49, 50, 51].

Here we present a high-quality genomic assembly and gene set of *H. incana*. As the first reference genome of a C_3_ plant with an exceptional rate of photosynthesis at high irradiance, we expect it to lay the foundation for studying photosynthetic light-use efficiency. First, we directly compare the photosynthetic rate of *H. incana* at high irradiance to that of the Brassicaceae species *B. rapa*, *B. nigra*, and *A. thaliana* to affirm its high light-use efficiency. Second, we characterize how the *H. incana* genome differs from that of other members of the Brassicaceae family, specifically focusing on differences in numbers of gene copies. Finally, we report whether such differences translate to differential expression of genes expected to mediate high light-use efficiency. Our work demonstrates how the assembly of *H. incana* serves as a valuable resource to elucidate the genetic basis of high photosynthetic performance and for studying the evolution of this trait in the Brassicaceae family.

## Results

### *Hirschfeldia incana* has an exceptionally high rate of photosynthesis

High net photosynthesis rates of more than 40 μmol · m^−2^·s^−1^ have been observed for *H. incana* in 1980 [31]. Net photosynthesis rates are those directly measured by gas-exchange measurement systems and reflect the net CO_2_ assimilation of plants while taking into account cellular respiration, a process which emits part of the total assimilated CO_2_ back into the atmosphere. We performed new measurements in order to compare the performance of *H. incana* with that of close relatives and the well-established model species *Arabidopsis thaliana*, grown under identical controlled environment conditions (Figure 1). Net CO_2_ assimilation rates did differ significantly between species (Additional File 2: Table S1). When leaves were adapted to the highest irradiance in our light response curve (2200 μmol · m^−2^·s^−1^), the highest net CO_2_ assimilation rates for *H. incana* accessions “Burgos” and “Nijmegen” were measured to be 50.8 and 46.6 μmol · m^−2^·s^−1^, respectively, much higher than the highest net CO_2_ assimilation rates measured for *B. nigra*, *B. rapa*, and *A. thaliana* (39.4, 38.7, and 32.3 μmol · m^−2^·s^−1^). The two *H. incana* accessions had the highest average net CO_2_ assimilation rates from 1100 μmol · m^−2^·s^−1^ irradiance, although only “Burgos” had a statistically significant higher rate than the other species (Additional File 2: Table S2). The same was found for gross photosynthesis rate (Additional File 2: Table S2), which is independent of the plant respiration rate (Additional File 2: Table S3).

**Figure 1:**
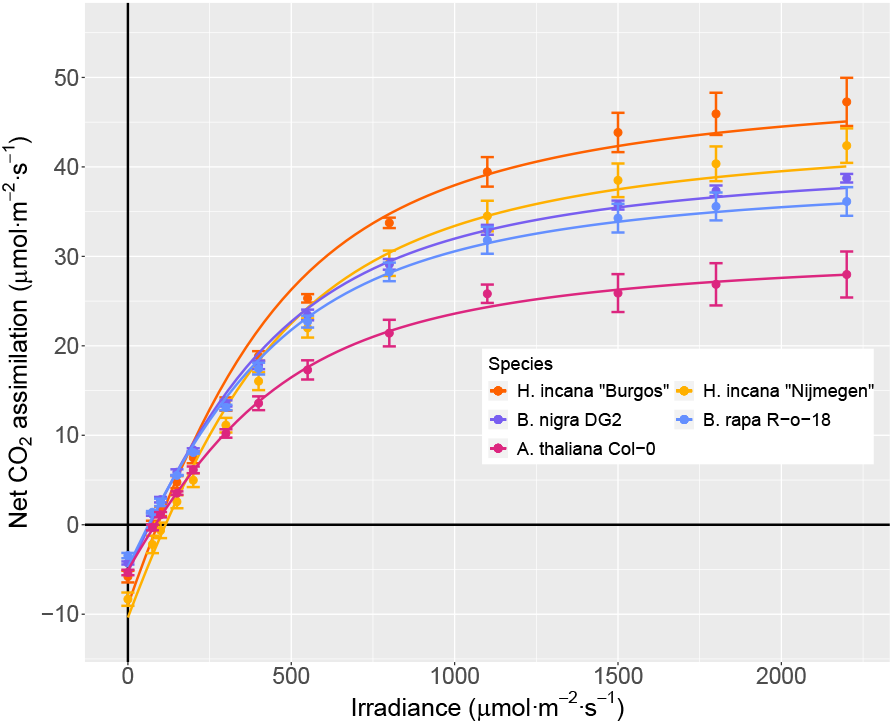
Two *H. incana* genotypes have a higher net CO_2_ assimilation at high irradiance than genotypes of close relatives. Light-response curves for *H. incana*, *B. rapa*, *B. nigra*, and *A. thaliana* accessions adapted to high levels of irradiance. Each point represents the average net CO_2_ assimilation value of three (*B. rapa*) or four leaves coming from independent plants. Error bars represent the standard error of means. The lines represent non-rectangular hyperbolas fitted on net assimilation rates for each species.

### A high-quality reference genome of *H. incana*

We assembled a high-quality reference genome of *H. incana* based on one genotype of the “Nijmegen” accession, that was inbred for six generations. Its haploid genome size was estimated to be 487 Mb, based on flow cytometry (Additional File 2: Table S4). This estimate is smaller than the previously reported genome size estimates of *B. rapa* (529 Mb) and *B. nigra* (632 Mb) [52]. Chromosome counts from root tip squashes showed 7 pairs of chromosomes (2n=14) (Additional File 1: Figure S1), consistent with previous reports [53, 29].

We generated DNA sequencing data consisting of 56 Gb of PacBio long reads (115-fold genome coverage, based on the genome size estimate), 46 Gb of 10X Genomics synthetic long reads (94-fold coverage, referred to as “10X” from now on for brevity), and 33 Gb of Illumina paired-end short reads (68-fold coverage). In addition, we generated 7.5 Gb of RNA sequencing data from leaf tissue for annotation purposes. Summary statistics and accession numbers can be found in Additional File 2: Table S5. A k-mer analysis of Illumina data resulted in a haploid genome size estimate of 325 Mb, with a low level of heterozygosity (1.2%).

Using a hybrid assembly strategy, we produced a nuclear genome assembly of 399 Mb of sequence in 384 scaffolds with an N50 of 5.1 Mb (Table 1). The assembly size is slightly larger than the genome size estimated from Illumina read k-mers (325 Mb), but smaller than the typical overestimate [54] based on flow cytometry (487 Mb). Besides the nuclear genome, we assembled the mitochondrial and chloroplast genome of *H. incana* into single sequences of 253 and 153 kb, and annotated the latter. The chloroplast assembly is typical for a Brassicaceae species, as it is nearly identical to chloroplast assemblies of *A. thaliana, B. rapa*, and *B. nigra* in terms of length and number of annotated genes (Additional File 2: Table S6).

**Table 1:** Genomic properties of assemblies generated of *H. incana* “Nijmegen” (this study), *B. rapa* Chiifu 401-42 [38] and *B. nigra* Ni100 [39].

|  | *H. incana* | *B. rapa* | *B. nigra* |
| --- | --- | --- | --- |
| Technologies | PacBio, 10X, Illumina paired-end | PacBio, BioNano, Hi-C, Illumina mate-pair | Nanopore, Hi-C, genetic mapping |
| Size (Mb) | 398.5 | 353.1 | 506 |
| # scaffolds | 384 | 1301 | 58 |
| N50 (Mb) | 5.1 | 4.4 | 60.8 |
| Gaps (kb) | 0.54 | 0.40 | 13 |
| GC-content (%) | 36.2 | 36.8 | 37.0 |
| Complete BUSCOs assembly (%) | 96.2 | 97.7 | 97.0 |
| Complete BUSCOs annotation (single/duplicated) (%) | 95.1 (80.2/14.9) | 97.2 (84.2/13.0) | 97.2 (81.9/15.3) |
| # protein-coding genes | 32,313 | 46,250 | 59,852 |
| # protein-coding transcripts | 38,706 | 46,250 | 59,852 |
| Repeat content (%) | 49.4 | 37.5 | 54.0 |
| Full-length LTR-RTs (%) | 25.3 | 29.2 | 41.8 |

The assembly is near-complete and structurally consistent with the underlying read data of *H. incana* “Nijmegen” (Additional File 2: Table S7). The high mapping rate of Illumina and 10X reads (>93%) suggest completeness, while the lower mapping rate of PacBio reads (81.5%) suggests some misassemblies or missing regions, likely repeats. The high mapping rate of RNA-seq reads (93.6%) again shows the gene space is near complete. We estimated the base-level error rate of the assembly to be 1 per 50 kb at most, based on variant calling using the mapped reads, resulting in 8,374 and 4,166 homozygous variants from the Illumina and 10X read alignments respectively.

We have annotated 32,313 gene models and 38,706 transcripts in the *H. incana* assembly (Table 1). This is a conservative annotation, based on filtering 64,546 initial gene models resulting from *ab initio*, protein alignment, and RNA-seq based predictions. Our filtering approach is more stringent than those used to generate the *B. rapa* and *B. nigra* annotations, which explains why we report a lower number of genes and transcripts for *H. incana* (Table 1) than for both *Brassica* species. Nevertheless, the annotation is expected to cover the large majority of the *H. incana* gene space. It contains 95.1% of 1,440 single-copy orthologs (BUSCOs) conserved in the Embryophyta plant clade, comparable to the percentages found for *B. rapa* and *B. nigra* (both 97.2%) (Table 1). The ratio of single to multiple copies is similar to that of *B. rapa* and *B. nigra* (Table 1), suggesting that the 14.9% of the BUSCOs present in multiple copies are true gene duplications shared by several species of the Brassiceae tribe. We additionally evaluated the completeness of the annotation by aligning proteins of *B. rapa* to the assembly and determining overlap between protein alignments and annotated genes. 30,552 out of the 37,387 protein alignments (81.7%) corroborate the annotation, as they completely or partially overlap with an annotated protein-coding gene. 2,570 (6.9%) of the protein alignments completely or partially overlap with an annotated repeat, suggesting that the aligned *B. rapa* proteins correspond to transposable elements. The remainder of the *B. rapa* proteins completely or partially overlap with gene models that were filtered (3,945 or 10.6%) or do not overlap with any annotated element at all (320 or 0.9%), indicating a small number of genes that are potentially missing from the annotation.

Based on these observations, we conclude that the *H. incana* assembly is mostly contiguous, correct, and complete, making it a solid foundation for comparative analyses with other Brassicaceae.

### The genome of *H. incana* extensively diversified from that of *B. rapa* and *B. nigra*

We utilized our assembly to explore the genomic divergence between *H. incana*, *B. rapa*, and *B. nigra*, all members of the same Brassiceae tribe. A substantial degree of divergence is expected between the three species due to different processes of post-polyploid diploidization, i.e. the process in which polyploid genomes get extensively rearranged as they return to a diploid state [55], following the ancient two-step genome triplication event shared by all Brassiceae [56, 36, 57]. Part of this divergence may have facilitated the evolution of the exceptional rate of photosynthesis at high irradiance of *H. incana*, if this trait was generated through natural selection.

We first recalibrated the time of divergence between *H. incana, B. rapa*, and *B. nigra* determined in earlier studies. A previous phylogenetic analysis based on four intergenic chloroplast regions suggested that *H. incana* is evolutionarily more closely related to *B. nigra* than to *B. rapa* and should be considered part of the “Nigra” clade [58] of the Brassiceae tribe. In contrast, a more recently constructed phylogeny of the Brassicaceae based on 113 nuclear genes implied that the *H. incana-B. nigra* split predates that of *H. incana-B. rapa* [59]. Our assembly suggests that the discrepancy between the two phylogenies is caused by the small difference in time between the speciation events of *H. incana-B. nigra* and *H. incana-B. rapa*, which we estimate to have happened respectively 10.35 and 11.55 million years ago (mya), based on the median rate of synonymous substitutions between their syntenic orthologs (*K_s_*) (Figure 2a).

**Figure 2:**
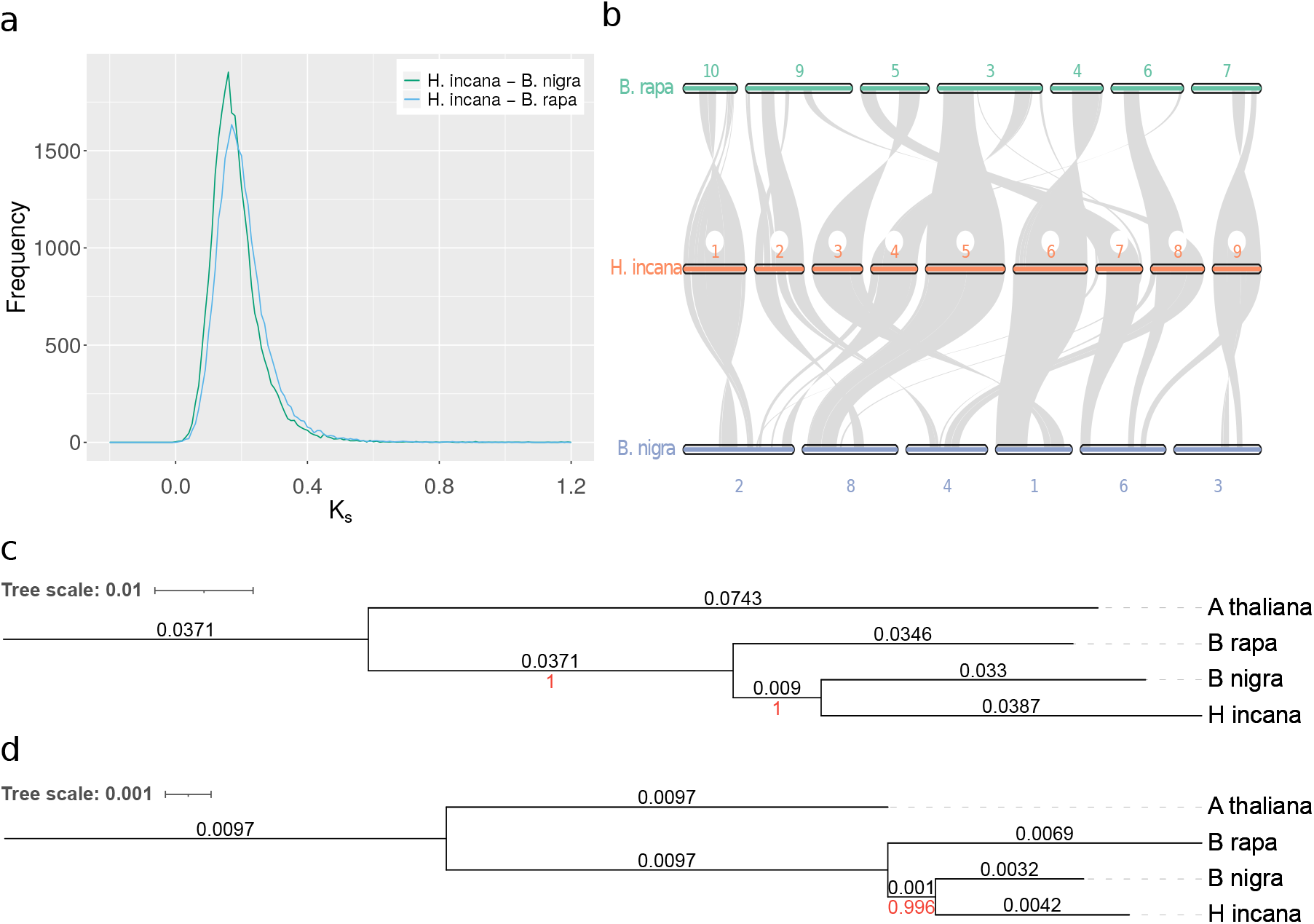
*H. incana* is evolutionary nearly equidistant from *B. rapa* and *B. nigra*, and has undergone several chromosomal rearrangements. (a) Distributions of the rates of synonymous substitutions between 25,127 and 26,137 orthologous gene pairs of *H. incana-B. rapa* and *H. incana-B. nigra*, respectively. Both distributions show a single clear peak corresponding to the speciation events. (b) Macrosynteny between the genomes of *H. incana, B. rapa*, and *B. nigra* genomes. Genes for which there exists a syntenic pair of orthologs in *H. incana-B. rapa* or *H. incana-B. nigra* are ordered along the scaffolds of *H. incana* and chromosomes of *B. rapa* and *B. nigra* based on sequence position. Pairs of syntenic orthologs are connected by gray lines. Only the nine largest scaffolds of *H. incana* (27% of the assembly) and chromosomes of *B. rapa* and *B. nigra* containing syntenic orthologs of genes in these scaffolds are shown for clarity. (c-d) Phylogenetic trees of *H. incana, B. rapa, B. nigra*, and *A. thaliana* (outgroup), based on nuclear (c) and chloroplast (d) genes. Branch lengths (black) and bootstrap values (red) are displayed above and below each branch, respectively.

We determined rearrangements between the genomes of *H. incana-B. rapa* and *H. incana-B. nigra* by comparing the order of syntenic orthologs between their assemblies. On a small scale, most genomic regions of *H. incana* are syntenic (not rearranged) to *B. rapa* and *B. nigra*, as 77.7% and 81.0% of the genes of *H. incana* could be clustered in collinear blocks containing a minimum of four orthologous pairs of *H. incana- B. rapa* and *H. incana-B. nigra*, respectively. Gene order is less conserved when comparing larger blocks, indicating several chromosomal rearrangements between the genomes of *H. incana* and the other two species (Figure 2b). For example, the two largest scaffolds of the *H. incana* assembly both contain inversions and/or translocations relative to their homologous chromosomes in *B. rapa* and *B. nigra* (Figure 2b). A similar pattern of rearrangements of small collinear blocks was observed between the genomes of *B. rapa* and *B. nigra* in past work [57].

A phylogenetic tree based on homologous nuclear genes corroborates the small difference in relatedness between *H. incana-B. nigra* and *H. incana-B. rapa* (Figure 2c). In contrast, *H. incana* is relatively more closely related to *B. nigra* than *B. rapa* according to a tree based on chloroplast genes (Figure 2d), suggesting the occurrence of at least one chloroplast capture event of unknown direction between *H. incana* and *B. nigra*. As we assume that the differences between the nuclear genomes are more reflective of the speciation events than those between the chloroplast ones, we propose that it is appropriate to consider *H. incana* as being evolutionarily nearly equally distant from both species, rather than assigning it to a specific clade.

We further examined genomic differentiation between the three species by comparing their transposable element (TE) content. The assembly of *H. incana* consists for 49.4% out of repetitive elements (Table 1), of which most are long terminal repeat retrotransposons (LTR-RTs) (25.3% of the genome). These numbers are consistent with previous work that investigated repeat content of the *H. incana* genome using genome skimming, which reported a repeat content of 46.5% and LTR-RT content of 31.6% [60]. We specifically focused our analyses on LTR-RTs, as LTR-RT expansion and contraction has been previously identified as a major driver of genomic differentiation between Brassiceae [61], even between different ecotypes of the same species [62]. The composition of LTR-RTs in the *H. incana* assembly differs from that of the *B. rapa* and *B. nigra* assembly, as it is dominated by Gypsy elements, consistent with earlier work [60], while Copia retrotransposons are dominant in the others (Figure 3a). Furthermore, the estimated insertion times of LTR-RTs varies between the three assemblies, as Gypsy and Copia elements in *H. incana* and *B. rapa* were predicted to have proliferated recently (< 1 mya) (Figure 3b-c), while Gypsy elements in *B. nigra* show a more varied distribution of insertion times (Figure 3b). A possible explanation of this shift could be that the *B. nigra* assembly was generated using longer reads than those used for the assemblies of *H. incana* and *B. rapa*, enabling it to capture a larger proportion of the centromeric regions, but we found no evidence that this introduced a bias towards longer insertion times of Gypsy elements (Figure 3b).

**Figure 3:**
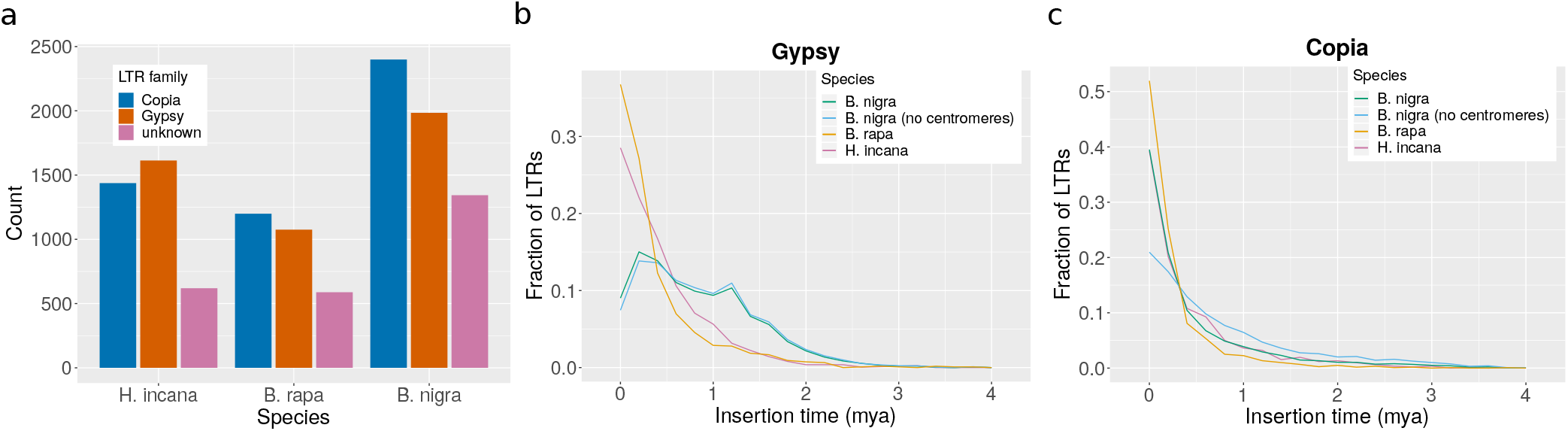
*Hirschfeldia incana* differs in Long Terminal Repeat Retrotransposon (LTR-RT) content from *B. rapa* and *B. nigra*. (a) Frequency distribution of LTR-RT families. LTR-RTs are classified as unknown if they contained elements of both Gypsy and Copia sequences and could thus not be reliably assigned to either of these families. (b) Frequency polygon (bin width = 0.2 mya) of the insertion times of Gypsy elements. (c) Frequency polygon (bin width = 0.2 mya) of insertion times of Copia elements.

Taken together, the breakdown of genomic synteny and divergence of LTR-RT content indicate that the genome of *H. incana* extensively diversified from that of *B. rapa* and *B. nigra* following their shared genome triplication event.

### Gene copy number variation may explain the high photosynthetic rate of *H. incana*

Genomic differentiation can result in species-specific gains and losses of genes, which may explain the differences in photosynthetic light-use efficiency between *H. incana*, *B. rapa*, and *B. nigra*. Given that the three species all share the same ancient genome triplication event [63, 57], it is reasonable to assume that most differences originated through differential retention of duplicated genes, particularly those located in genomic blocks showing evidence of extensive fractionation since that event [57]. We investigated gene copy number variation between the three species, by clustering their annotated protein-coding genes with those of five other Brassicaceae species within (*R. raphanistrum* and *R. sativus*) and outside (*A. arabicum*, *A. thaliana*, and *S. irio*) the Brassiceae tribe into homology groups. The inclusion of *A. thaliana* allowed us to use its extensive genomic resources to functionally annotate the genes of other species. The other four species were included to put the analysis in a broader phylogenetic context. *A. arabicum* is part of the Aethionema tribe which diverged from the core group of the Brassicaceae family, thus allowing us to identify highly conserved genes. *S. irio* is part of a different tribe than *A. thaliana* (Sisymbrieae), that is more closely related to the Brassiceae tribe [59], but did not undergo the ancient genome triplication. *R. raphanistrum* and *R. sativus* are part of the *Raphanistrum* genus within the Brassiceae tribe and thus represent another set of species that underwent the genome triplication shared by the whole tribe.

Our analysis resulted in 20,331 groups containing at least one *H. incana* gene (Additional File 2: Table S8). The composition of the homology groups agrees with the currently established phylogeny of the Brassicaceae [59], as groups containing *H. incana* genes share the fewest genes with *A. arabicum* (58.2%) and most genes with species part of the Brassiceae tribe (86.3-95.6%). *H. incana* has a low fraction of species-specific homology groups (3.4%) compared to the seven other species, which can be attributed to the stringent filtering of the predicted gene models.

We focused on a subset of 15,097 groups containing at least one gene of *A. thaliana* and one of *H. incana*, as these could be extensively annotated through the transfer of Gene Ontology (GO) terms from *A. thaliana* genes to their respective groups. According to the expectation that most genes quickly return to single-copy status following a whole genome duplication event [64], 70.2% of these groups contain a single gene of both *A. thaliana* and *H. incana*. Focusing on groups containing *A. thaliana* genes involved in photosynthesis (260 in total, Additional File 2: Table S9), most contain a higher number of genes of *H. incana*, *B. rapa*, and *B. nigra*, compared to *A. thaliana* (Figure 4), consistent with the relatively higher photosynthetic light-use efficiency of the latter three (Figure 1). However, despite *H. incana* having the highest efficiency of all four species, most groups contain fewer or the same number of copies of this species, relative to *B. rapa* and *B. nigra*. This is not a result of our conservative filtering approach, as we explicitly retained putative photosynthesis-related genes during our filtering procedure (Additional File 1: Supplementary Methods).

**Figure 4:**
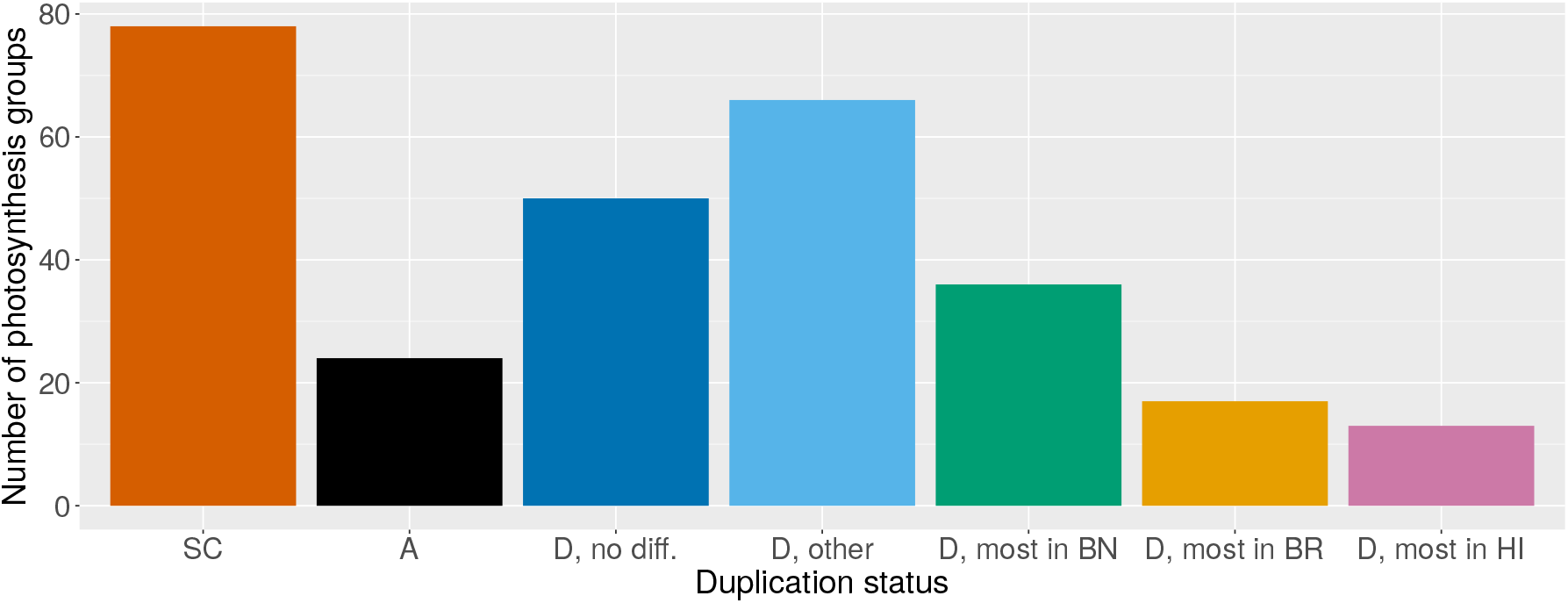
*H. incana* retained fewer duplicated copies of photosynthesis-associated genes than *B. rapa* and *B. nigra*. Bars show counts of homology groups containing genes associated with photosynthesis with different distributions of copy numbers in the four species (260 groups in total). Groups are classified as SC if they contain a single copy of all, A if they contain the highest number of copies in *A. thaliana*, and D if they contain a higher number of copies of *H. incana*, *B. rapa*, and *B. nigra* than of *A. thaliana*. If there is no difference in the number of copies of *H. incana*, *B. rapa*, and *B. nigra*, groups within the D category are classified as no diff. Otherwise, groups are labeled based on whether multiple species contain the highest number of copies (other) or whether there is a single one (*B. nigra* = most in BN, *B. rapa* = most in BR, *H. incana* = most in HI).

Besides photosynthesis-related genes, we also analysed CNVs of a more general set. 2,302 homology groups contain genes present at higher copy numbers in *H. incana*, *B. rapa*, and *B. nigra*, compared to *A. thaliana* (Additional File 2: Table S10), and these groups are significantly enriched for 81 GO terms (Additional File 2: Table S11). Overrepresented genes include those involved in regulation (e.g. of transcription) and those annotated with GO terms associated with interaction with the environment (e.g. “response to cadmium ion”), of which “response to light stimulus” is particularly intriguing, given that the increased photosynthetic light-use efficiency of *H. incana*, *B. rapa*, and *B. nigra*, relative to *A. thaliana*, is particularly pronounced at high levels of irradiance (Figure 1). Genes within the expanded groups associated with this term that could likely explain this phenomenon are those directly involved in photosynthetic processes, which include several light-harvesting complex genes (Additional File 2: Table S12).

As CNV can considerably affect gene expression levels [65], we hypothesized that retained copy number expansions of photosynthesis and high light response-related genes in *H. incana*, *B. rapa* and *B. nigra* may aid in the high photosynthetic capacities found in this study (Figure 1). We therefore measured gene expression levels of twelve genes for which there is inter-species CNV in two contrasting light conditions (200 μmol · m^−2^·s^−1^ and 1500 μmol · m^−2^·s^−1^) (Additional File 2: Table S13). We selected genes with GO annotations related to either photosynthesis or high light responsiveness or with a function associated with these processes described in literature. For eight of these twelve genes we find significantly higher gene expression levels at higher gene copy numbers while for the remaining four genes no such correlation is observed (Figure 5). This suggests that the increased copy numbers of photosynthesis and high light response-related genes in *H. incana*, *B. rapa* and *B. nigra*, relative to the *A. thaliana*, may indeed contribute to their high photosynthetic efficiency, although this effect appears to not be specific to a particular species or level of irradiance.

**Figure 5:**
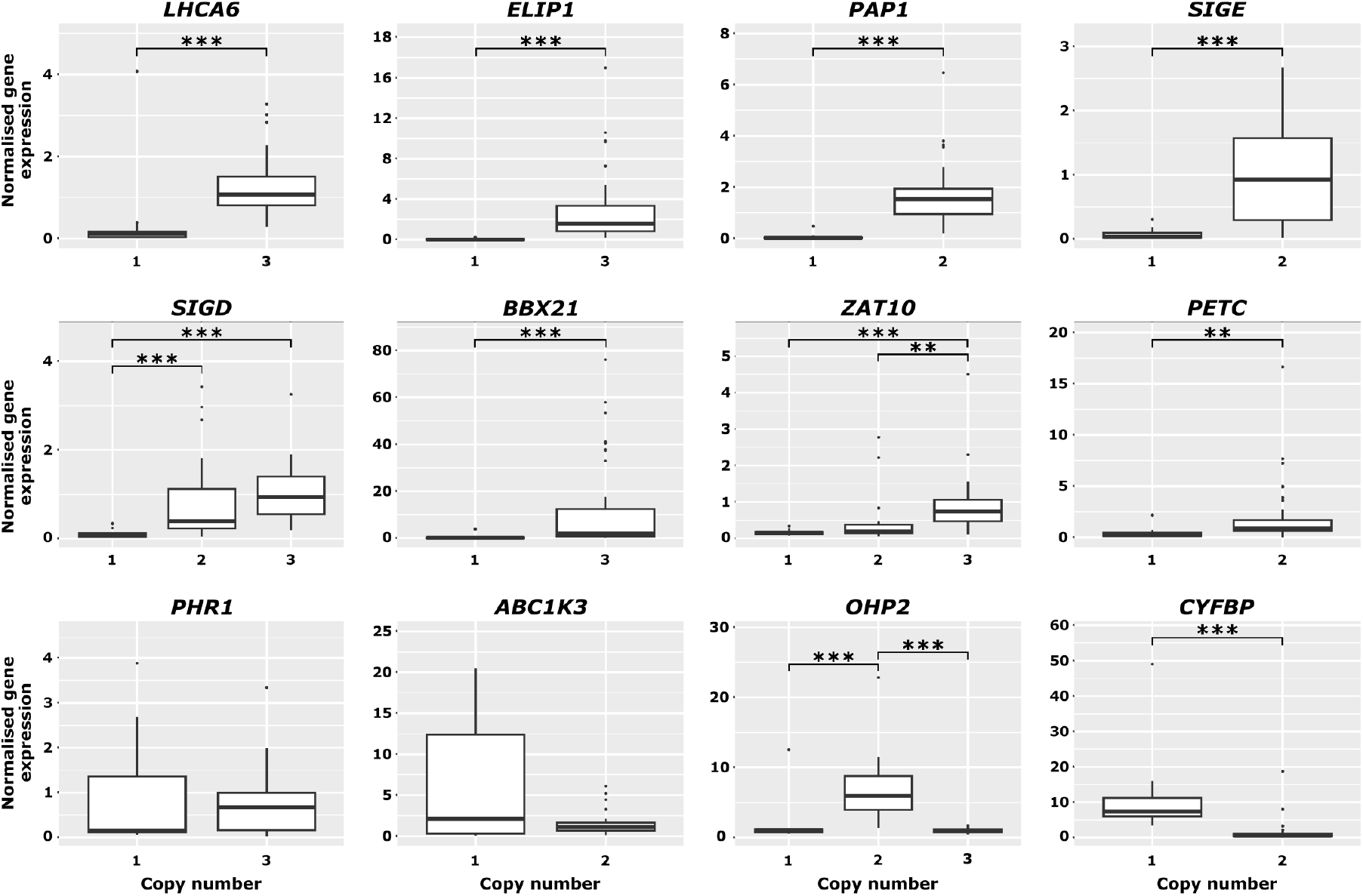
Copy numbers of photosynthesis-associated genes correlate with expression level. Boxplots depict gene expression levels of *A. thaliana, B. rapa, B. nigra* and *H. incana* grown in 200 μmolm^−2^ s^−1^ and 1500 μmolm^−2^ s^−1^. Gene expression levels were normalized against *H. incana* grown at 200 μmol m^−2^ s^−1^ and subsequently grouped per gene copy number. Titles of graphs indicate gene names based on the *A. thaliana* gene nomenclature. *p < 0.05; **p < 0.01; ***p < 0.001.

## Discussion

In this study, we generated a high-quality reference genome of *H. incana* to establish this species as a model for exceptional photosynthetic light-use efficiency at high irradiance. We find substantial differences in light-use efficiency, genomic structure, and gene content between *H. incana* and its close relatives. We discuss these results in terms of how they contributed to the evolution of the remarkable phenotype of *H. incana*.

We have validated and extended results showing the exceptional photosynthetic light-use efficiency at high irradiance of *H. incana*. Our measurements imply that the rates of photosynthesis of *H. incana* are higher than those of the C_4_ crop maize [32, 33] and remarkably, up to 50% and 100% higher than those of key cereal crop species with similar C_3_ photosynthetic metabolism, such as wheat [15] and rice [17]. Furthermore, these rates are higher than those of closely related Brassicaceae species *B. rapa*, *B. nigra*, and the more distantly related *A. thaliana*. We confirm that the positive correlation between net CO_2_ assimilation rates and chlorophyll content reported for many Brassicaceae species [66, 67] applies to *H. incana*, *B. nigra* and *B. rapa* too.

The photosynthetic rates measured for *H. incana* are higher than those reported in previous work [31], as are the rates measured in *B. rapa* [68, 69]. While we were unable to find previously published data for *B. nigra* and *A. thaliana* grown under high light, the measured rates of both species are higher than expected compared to the performance of C_3_ crops resp. previous studies of *A. thaliana* conducted under low irradiance conditions. Although the rates presented in this study were obtained from plants grown in controlled, favourable conditions and thus are likely to be an overestimation of rates in natural environments, the magnitude of the differences suggests that *H. incana* can provide essential information for the improvement of photosynthetic light-use efficiency in crops.

The high-quality reference genome of *H. incana* generated in this study provides the means to elucidate the genetic basis of this plant’s exceptional rate of photosynthesis and how it evolved in this species. We estimate that *H. incana* diverged 11.55 and 10.35 mya from *B. rapa* and *B. nigra*, respectively, consistent with an earlier study that used a smaller set of nuclear genes [59]. These time points are close to the reported time at which *B. rapa* and *B. nigra* (11.5 mya) [39] diverged from each other and the time at which the whole Brassicaceae family underwent a rapid radiation event [70]. It has been proposed that this event was mediated by the expansion of grass-dominated ecosystems in the native region of the Brassicaceae family at that time, which created newly opened habitats that favoured rapid diversification [70]. This expansion in turn, is thought to have been driven by decreasing atmospheric CO_2_ levels and increasing temperatures, which favoured the displacement of the then dominant C_3_ plants by C_4_ grasses [71]. We argue that these climatic changes drove the evolution of the highly efficient rate of photosynthesis observed in *H. incana* as well, analogous to how the C_4_ photosynthesis pathway evolved as an adaptation to low CO_2_ levels and drought.

Our analyses suggest that the genome of *H. incana* extensively differentiated from those of *B. rapa* and *B. nigra* since their time of divergence through large genomic arrangements and differences in LTR-RT content. We hypothesize that the latter observation is caused in part due to Gypsy elements being less efficiently purged from the genome of *B. nigra* than from those of *H. incana* and *B. rapa*. An increased rate of LTR-RT removal, based on the ratio of solo LTRs to intact LTR-RTs, has also been observed in *B. rapa* relative to *B. oleracea* and it was speculated that this is caused by the increased rate of genetic recombination in the former [72]. Given that a similar negative correlation between local recombination rate and LTR-RT content was found in rice [73] and soybean [74], the differences in predicted insertion times of Gypsy elements in *H. incana, B. rapa*, and *B. nigra* observed in this study may thus reflect different rates of genetic recombination in the three species. While it has been suggested that changes in recombination rate can be adaptive, there is little empirical evidence that supports this [75]. It would therefore be interesting to directly measure genomewide rates of recombination of *H. incana*, *B. rapa*, and *B. nigra* and explore whether these are linked to their rate of photosynthesis.

Further comparative analyses between the genomes of *H. incana*, *B. rapa*, *B. nigra*, and *A. thaliana* revealed numerous species-specific gains and losses of genes. For dosage-sensitive genes, such as those involved in transcriptional regulation, differences may not necessarily reflect adaptive selection. This category of genes was found to be consistently retained in multiple copies following polyploidy events across the Brassicaceae [76] and a wide group of angiosperms [64], which is hypothesized to be due to dosage constraints [77]. Differences in copy number of such genes may thus reflect different rates of relaxation of dosage balance constraints and subsequent loss of duplicates through time, which is a neutral process. On the other hand, there is reason to believe that gene duplications contributed to the evolution of the high light-use efficiency of *H. incana*. Gene duplications have been identified as important drivers of plant evolution and inter-species CNVs are often enriched for adaptive evolutionary traits [78, 79, 80]. In support of this hypothesis, genes duplicated in *H. incana*, *B. rapa*, and/or *B. nigra*, relative to *A. thaliana*, are enriched for the GO term “response to light stimulus”. Moreover, the expression levels of eight out of twelve duplicated genes associated with photosynthesis selected for RT-qPCR analysis correlate with gene copy number, independent of the level of irradiance.

In this context, it is striking that *H. incana* overall contains fewer photosynthesis-related genes than *B. rapa* and *B. nigra*. This points towards an alternative scenario in which adaptation of *H. incana* to high levels of irradiance occurred through regulation of expression of one copy of the photosynthesis-related genes, which relaxed selection on duplicate retention or even encouraged duplicate loss. Another explanation is that *H. incana* did not adapt through changes in core photosynthetic genes, but rather through changes in other traits, such as mesophyll conductance to CO_2_. To elucidate the exact genetic mechanisms underlying the high light-use efficiency of *H. incana*, a natural follow-up to this study is to perform comparative transcriptomic analyses of leaves of *H. incana*, *B. rapa*, and *B. nigra* under a range of different levels of irradiance and at different developmental stages. Genes that show copy number variation and are differentially expressed between *H. incana* and the latter two species would then be prime candidates to further test for potential causality through e.g. knock-out mutant analysis. As previous work has shown that it is possible to cross distantly related Brassicaceae species [81], a useful approach to further pinpoint the causal genes is to establish a genetic mapping population between *H. incana* and a Brassicaceae species with regular light-use efficiency and perform quantitative trait locus analyses of photosynthetic traits segregating within the population.

## Conclusions

*H. incana* has an exceptional rate of photosynthesis at high irradiance. We generated a near-complete reference genome of this species and found evidence suggesting that its exceptional rate evolved through differential retention of duplicated genes. Taken together, our results illustrate the promise of *H. incana* as a model organism for high photosynthetic light-use efficiency and we expect the reference genome generated in this study to be a valuable resource for improving this efficiency in crop cultivars.

## Materials and Methods

### Plant material

*Hirschfeldia incana* accessions “Nijmegen” and “Burgos” were used. “Nijmegen” is an inbred line (> six rounds of inbreeding) originally collected in Nijmegen, The Netherlands. Seeds of “Burgos” were originally collected at Burgos, Spain. Furthermore, *Brassica nigra* accession “DG2”, sampled from a natural population at Wageningen, The Netherlands, the *Brassica rapa* inbred line “R-o-18” [82, 83], and the *Arabidopsis thaliana* Col-0 accession were used.

### Measurements of photosynthesis rates

Seeds of *H. incana* “Nijmegen”, *H. incana* “Burgos”, *B. rapa* R-o-18, *B. nigra* “DG2”, and *A. thaliana* Col-0 were sown in 3 L pots filled with a peat-based potting mixture. Plants were grown in a climate chamber with a photoperiod of 12 hours and day and night temperatures of 23 and 20 °C, respectively. Humidity and CO_2_ levels were set at 70% and 400 ppm. The chamber was equipped with high-output LED light modules (VYPR2p, Fluence by OSRAM). Plants were watered daily with a custom nutrient solution (0.6 mm NH_4_^+^, 3.6 mm K^+^, 2 mm Ca^2+^, 0.91 mm Mg^2+^, 6.2 mm NO_3_^−^, 1.66 mm SO_4_^2^, 0.5 mm P, 35 μm Fe^3+^, 8 μm Mn^2+^, 5 μm Zn^2+^, 20 μm B, 0.5 μm Cu^2+^, 0.5 μm Mo^4+^). The seeds were germinated at an irradiance of 300 μmol · m^−2^·s^−1^, and the same irradiance was maintained to let seedlings establish. On day 14, 21, and 25 after sowing, the irradiance was raised to 600, 1200, and 1800 μmol · m^−2^·s^−1^, respectively.

The photosynthetic metabolism of young, fully expanded leaves developed under 1800 μmol · m^−2^·s^−1^ of light was measured with a LI-COR 6400xt portable photosynthesis system (LI-COR Biosciences) equipped with a 2 cm^2^ fluorescence chamber head. “Rapid” descending light-response curves were measured between 30 and 35 days after sowing to accommodate differences in growth rates of the different species on one leaf from four *H. incana* “Nijmegen”, *H. incana* “Burgos”, *B. nigra* “DG2”, and *A. thaliana* Col-0 plants, and three *B. rapa* R-o-18 plants. The net assimilation rates of the plants were measured at thirteen different levels of irradiance ranging from 2200 to 0 μmol · m^−2^·s^−1^. During measurements, leaf temperature was kept constant at 25 °C and reference CO_2_ concentration was kept at 400 μmol · mol^−1^. Water in the reference air flux was regulated in order to achieve vapour-pressure deficit values comprised between 0.8 and 1.2 kPa.

Light response curve parameters (*A_max_*: net CO_2_ assimilation at saturating irradiance, *φ*: apparent quantum yield of CO_2_ assimilation, *R_d_*: daytime dark respiration rate, and *θ*: curve convexity) were estimated for each species trough nonlinear least squares regression of a non-rectangular hyperbola [84] with the R package “photosynthesis” (version 2.0.0) [85]. An indication of gross assimilation rates for each species was subsequently generated by adding the daytime dark respiration rate (*R_d_*) estimated for each species to the species’ net assimilation rates.

Differences in net and gross assimilation rates were tested at each light level of the light-response curve with a oneway ANOVA on the “genotype” experimental factor. Pairwise comparisons between the assimilation rates of the different genotypes at each light level were subsequently performed and tested with the Tukey-Kramer extension of Tukey’s range test. The *p*-value threshold for statistical significance was set at *α* = 0.05.

### Flow cytometry

Leaf samples of the *H. incana* genotypes “Burgos” and “Nijmegen” and *A. thaliana* ecotype Col-0 were analysed for nuclear DNA content by flow cytometry (Plant Cytometry Services B.V., Didam, the Netherlands). Seven, three and five biological replicates were measured over separate rounds of analysis for *H. incana* “Nijmegen” *H. incana* “Burgos”, and *A. thaliana* Col-0, respectively. Nuclei were extracted from leaf samples following the method by [86], and stained with 4’,6-diamidino-2-phenylindole (DAPI). The DNA content of nuclei relative to that of the reference species *Monstera deliciosa* was determined on a CyFlow Ploidy Analyser machine (Sysmex Corporation, Kobe, Japan). A haploid flow cytometry estimate of 157 Mb was used for *A. thaliana*, resulting from comparisons of nuclear DNA content of this species and other model organisms [87]. Haploid genome size estimates for the *H. incana* genotypes were obtained by multiplying the *H. incana-to-M. deliciosa* ratio by the haploid *A. thaliana* estimate and dividing this product by the average *A. thaliana*-to-*M. deliciosa* ratio.

### Chromosome counting

Root tips (approximately 1 cm long) were collected from young, fast-growing rootlets of multiple *H. incana* “Nijmegen” plants and pre-treated for 3 h at room temperature with a 0.2 mm 8-hydroxyquinoline solution. After pre-treatment, the 8-hydroxyquinoline solution was replaced with freshly prepared Carnoy fixative (1:3 (v/v) acetic acid - ethanol solution) and maintained at room temperature for half a day. Root tips were then rinsed with 70% ethanol for three times to remove remaining fixative and stored in 70% ethanol at 4 °C until further use. Prior to slide preparation, root tips were rinsed twice in MQ water before adding 1:1 solution of a pectolytic enzymatic digestion solution (1% Cellulase from *Trichoderma*, 1% Cytohelicase from *Helix Promatia*, 1% Pecolyase from *Aspergillus japonicus*) and 10 mm citric buffer. After one hour incubation at 37 °C, the enzymatic digestion solution was replaced by MQ water. The digested root tips were spread in 45% acetic acid over microscopy slides on a hot plate set at 45 °C, cells were fixed with freshly prepared Carnoy fixative, dried, and stained with 4’,6-diamidino-2-phenylindole (DAPI) dissolved in Vectashield mounting medium (Vector Laboratories Inc., Burlingame, US). Slides were imaged with an Axio Imager.Z2 fluorescence optical microscope coupled with an Axiocam 506 microscope camera (Carl Zeiss AG, Oberkochen, Germany) at 63x magnification. Chromosome numbers were counted in metaphase mitotic cells and averaged to obtain the reported number.

### DNA and RNA isolation

Genomic DNA was extracted from *H. incana* “Nijmegen” samples using a protocol modified from [88]. The modifications consisted of adding 300 μL *β*-mercaptoethanol to the extraction buffer just before use. We added 0.7% isopropanol to the supernatant instead of 10 m LiCl and then divided the total volume into 1 mL aliquots for subsequent extractions. The pellet was dissolved in 500 μL of SSTE which was preheated to 50 °C before use. The final pellets were dissolved in 50 μL Milli-Q and then pooled at the end of the extraction process. DNA used for Illumina and 10X Genomics sequencing was extracted from flower material, while leaf material was used for the PacBio sequencing, all originating from the same plant.

Total RNA was extracted from leaf material of *H. incana* “Nijmegen” from a different plant than the one used for the DNA isolations with the Direct-zol RNA mini-prep kit (Zymo Research) according to the company’s instructions and then subjected to a DNAse (Promega) treatment at 37 °C for one hour.

### Generation of sequencing data

Sequencing of total-cellular DNA of *H. incana* “Nijmegen” was performed by GenomeScan B.V., Leiden. A total of seven SMRT cells were used for sequencing on the Pacific Biosciences Sequel platform. Short read Illumina and 10X Genomics libraries with an insert size of approximately 500-700 bp were prepared with the NEBNext Ultra DNA Library Prep kit for Illumina and 10X Genomics ChromiumTM Genome v1 kit, respectively. These libraries were sequenced using the Illumina X10 platform (2 × 151 bp). RNA paired-end sequencing libraries with an average insert size of 254 bp were prepared using the Illumina TruSeq RNA sample prep kit with polyA mRNA selection and sequenced using the Illumina HiSeq 2500 platform (2 × 125 bp).

### k-mer analysis

A histogram of k-mer frequencies of Illumina reads predicted to be of nuclear origin (see Additional File 1: Supplementary Methods) was generated using Jellyfish (v2.2.6) [89], using a k-mer length of 21. The resulting histogram was provided as input to Genomescope (v1.0.0) [90] to estimate genome size and heterozygosity.

### Genomic assembly and annotation

The genome assembly and annotation process is more extensively described in Additional File 1: Supplementary Methods. In short, we generated an initial assembly based on the PacBio data only with Canu [91] and used it to bin the PacBio, 10X, and Illumina reads according to whether they originated from nuclear, organellar, or contaminant DNA. The bins were used to separately assemble the nuclear and organellar genomes, yielding a nuclear assembly consisting of hundreds of contigs and mitochondrial and chloroplast assemblies that were both represented by a single sequence. Nuclear contigs representing alternative haplotypes were removed using purge_dups [92], after which ARKS [93] was used to scaffold the remaining contigs using the 10X data. Scaffolds were polished using Arrow (https://github.com/PacificBiosciences/gcpp) and Freebayes [94], followed by a manual filtering step to obtain the final nuclear assembly.

Repeats in the assembly were masked using Repeat-Masker [95] in combination with RepeatModeler2 [96] before starting the annotation procedure. Nuclear genes were annotated by using EvidenceModeler [97] to generate consensus models of ab initio gene predictions, alignments of proteins from closely and distantly related plant species, and transcripts assembled from RNA-seq data. These models were manually filtered to obtain a final set of protein-coding genes.

### Used datasets for comparative genome analyses

We mainly focused comparative genome analyses on *H. incana*, *B. nigra*, and *B. rapa*, three species of the Brassiceae tribe of which all members underwent an ancient genome triplication [56, 36]. For comparative gene analyses, we extended this group with the Brassicaceae species *Arabidopsis thaliana*, *Aethionema arabicum*, *Sisymbrium irio*, *Raphanus raphanistrum*, and *Raphanus sativus*. The latter two *Raphanus* species are also part of the Brassiceae tribe. Version numbers and locations of all genomes are listed in Additional File 2: Table S14.

### Analysis of pairwise gene synteny and long terminal repeat retrotransposons (LTR-RTs) in *H. incana, B. rapa*, and *B. nigra*

Analyses of pairwise gene synteny between scaffolds of *H. incana* and chromosomes of *B. rapa* and *B. nigra* were performed using the JCVI library (https://github.com/tanghaibao/jcvi) (v1.0.5) in Python. Orthologs were identified through all-vs-all alignment of genes with LAST [98], retaining reciprocal best hits only (C-score of at least 0.99). Hits were filtered for tandem duplicates (hits located within 10 genes from each other) and chained using the Python implementation of MCScan [99] to obtain collinear blocks containing at least four pairs of syntenic genes. Visualizations of collinearity between genomic assemblies were generated using custom scripts of JCVI.

*K_s_* values of syntenic gene pairs were computed using the ks module of JCVI. Protein sequences of pairs were aligned against each other using MUSCLE (v3.8.1) [100], after which PAL2NAL (v14) [101] was used to convert protein alignments to nucleotide ones. *K_s_* values for each pair were computed from the nucleotide alignments using the method of [102] implemented in PAML [103] (v4.9). Times of divergence between species were estimated by dividing the median of the distributions of their *K_s_* values by the rate of 8.22 × 10^−9^ synonymous substitutions per year that was established for Brassicaceae species based on extrapolation from the ancient triplication event [104].

Putative LTR-RTs were identified using LTRharvest (v1.6.1) [105] and LTR_finder (v1.1) [106], after which LTR_retriever (v2.9.0) [107] was run with default parameters to filter and combine the output of both tools into a high confidence set. LTR_retriever was also used to provide estimates of the insertion time of each LTR-RT. Parameters of LTRharvest and LTR_finder were set as recommended in the LTR_retriever documentation. Centromeric regions of the *B. nigra* assembly were obtained from Table S21 of the manuscript describing the assembly [39].

### Phylogenetic analysis of *H. incana, B. rapa*, and *B. nigra*

The longest isoforms of the nuclear genes of *H. incana*, *B. rapa*, *B. nigra*, and *A. thaliana* (outgroup) were provided to Orthofinder (version 2.3.11) [108] to generate phylogenetic species trees. Orthofinder was run using the multiple sequence alignment (MSA) workflow with default parameters. The same analysis was performed using chloroplast genes. Trees were visualized using iTOL (version 6.3) [109].

### Comparative gene ontology analysis of eight Brassicaceae species

The longest isoforms of the genes of all eight Brassicaceae species described in the section “Used datasets for comparative genome analyses” were extracted using AGAT (version 0.2.3) (https://github.com/NBISweden/AGAT) and clustered into homology groups using the “group” function of Pantools version 2 [110] with a relaxation parameter of 4. Groups were assigned GO slim terms of their associated *A. thaliana* genes (obtained from https://www.arabidopsis.org/download_files/GO_and_PO_Annotations/Gene_Ontology_Annotations/TAIR_GO_slim_categories.txt (last updated on 2020-07-01)) and GO terms assigned to protein domains of associated *H. incana*, *B. rapa*, and *B. nigra* genes using InterProScan (version 5.45-72.0) [111] (ran using the Pfam and Panther databases only). GO term enrichment tests were performed using the Fisher Exact test and the Benjamini-Hochberg method for multiple testing correction [112]. *A. thaliana* genes were considered to be involved in photosynthesis, if they fulfilled one of the following conditions:

- Annotated with one of the following GO terms: “photosynthesis”, “electron transporter, transferring electrons within the cyclic electron transport pathway of photosynthesis activity”, or “electron transporter, transferring electrons within the noncyclic electron transport pathway of photosynthesis activity”;
- Included in the KEGG pathways ath00195 (Photosynthesis), ath00710 (Carbon fixation in photosynthetic organisms), and ath00196 (Photosynthesis - Antenna Proteins);
- Protein products have been assigned the keyword “Photosynthesis” in the Swiss-Prot database.

The same criteria were used to retain photosynthesis-related genes of *H. incana* while filtering the gene annotation of the assembly (see Additional File 1: Supplementary Methods).

### Analysis of gene expression under high and low irradiance

Seeds of *H. incana* “Nijmegen”, *B. rapa* “R-o-18”, and *B. nigra* “DG2” were vapor-phase sterilised for 7 h following a previous protocol [113], and germinated on 1/2 MS medium for five days at 25 °C and 100 μmol · m^−2^·s^−1^ with a photoperiod of 16 h. Sixteen seedlings per species were then transferred to 2 L pots filled with a peat-based potting mixture.

Plants were grown in a climate chamber with a photoperiod of 12 hours and day and night temperatures of 23 and 20 °C, respectively. Humidity and CO_2_ levels were set at 70% and 400 ppm. The chamber was equipped with high-output LED light modules (VYPR2p, Fluence by OSRAM, Austin, USA). Eight plants per species were assigned to a high light (HL) treatment of 1500 μmol · m^−2^·s^−1^ and the remaining eight to a low light (LL) treatment of 200 μmol · m^−2^·s^−1^.

Plants assigned to the HL treatment were gradually adapted to target irradiance by starting at 200 and raising irradiance to 800 and 1500 μmol · m^−2^·s^−1^, 9 and 13 days after sowing, respectively. Pots were watered daily with the same custom nutrient solution as the one used in the measurements of photosynthesis rates and twice a day when irradiance was raised to 1500 μmol · m^−2^ ·s^−1^. Plants assigned to the LL treatment were always grown at 200 μmol · m^−2^·s^−1^ and watered daily with the same custom nutrient solution.

Twenty days after sowing, one young fully adapted leaf from each plant was selected, excised, and snap-frozen in liquid nitrogen. Due to space limitations, a repetition of this experiment had to be subsequently performed employing the same procedures to allow the inclusion of *A. thaliana* Col-0 plants in this experiment. In order to have a reference between both repetitions, *A. thaliana* plants were grown together with *H. incana* “Nijmegen” plants.

Leaf samples were crushed with a mortar and pestle cooled with liquid nitrogen and further homogenised with glass beads for 2 min at 30 Hz in a MM300 Mixer Mill (Retsch GmbH, Haan, Germany). Total RNA was extracted with the RNeasy Plant Mini Kit (QIAGEN N.V., Venlo, The Netherlands) according to manufacturer’s instructions and then subjected to a RQ1 DNAse treatment (Promega Corporation, Madison, U.S.) at 37 °C for 30 minutes. We validated the total removal of DNA by means of a no reverse transcriptase PCR reaction on all RNA samples. The RNA quality was assessed for purity (A260/A280) with a Nanodrop 2000 spectrophotometer (Thermo Fisher Scientific Inc., Waltham, U.S.) and for possible RNA degradation by means of a visual inspection of the RNA on a 1% agarose gel. cDNA was then synthesized from 2 μg total RNA (measured by spectrophotometer) with the SensiFAST™ cDNA Synthesis Kit (Meridian Bioscience, Cincinnati, U.S.) according to manufacturer’s instructions.

To examine the expression of photosynthesis-related genes (Additional File 2: Table S13), species-specific RT-qPCR primers were designed with the following criteria: the primer pair had to bind to all copies of a particular gene in one species, the PCR fragment size had to range between 80 and 120 bp, the maximum difference in melting temperature between primers of the same pair had to be 0.5 °C, and overall melting temperatures had to be comprised between 58 and 62 °C. Primers were designed to target a region of the gene as similar as possible region in all species. RT-qPCR reactions were performed with SYBR green on a CFX96 Real-Time PCR Detection System (Bio-Rad Laboratories Inc., Hercules, U.S.). The efficiency of each designed primer set was assessed by means of a standard curve, and only primer sets with efficiencies ranging between 90% and 110% were used. All primer sequences can be found in Table S18.

Gene expression was first normalized to the reference genes *ACT2, PGK, UBQ7* and *APR* [114, 115] using the delta-Ct (dCt) method [116]. Then, in order to compare the relative gene expression levels between the two experiments, one with *A. thaliana* and *H. incana* and the second with *H. incana*, *B. rapa* and *B. nigra*, we normalized the data against *H. incana* LL samples. This was done by calculating the average dCt of a respective gene in each of the two sets of *H. incana* samples grown in LL, and subsequently dividing the dCt of all other samples by the average of the *H.incana* LL samples grown in the same experiment. Normalized gene expression values were then calculated as 2^−dCt^. For the statistical analysis we log-transformed (log10) the normalized gene expression values and tested the significance with a oneway ANOVA. A post-hoc test was subsequently performed with Tukey’s range test, with the significance threshold set at *α* = 0.05.

## Availability of data and material

Raw sequencing data of *Hirschfeldia incana* can be found on the repository of the National Center for Biotechnology Information (NCBI) (BioProject ID: PRJNA612790). The assembly of *Hirschfeldia incana* has been deposited at DDBJ/ENA/GenBank under the accession JABCMI000000000. The version described in this manuscript is version JABCMI010000000.

## Authors’ contributions

JH and MGMA initiated the research on the genomic basis of high photosynthesis of *H. incana*. MGMA generated the inbred line of *H. incana* “Nijmegen”. FG performed measurements of photosynthesis rates, flow cytometry experiments, and subsequent analyses of experimental data. VC performed the gene expression experiment. VC and RB analysed the results of the gene expression experiment. RB and FB performed DNA and RNA. RYW and SS generated strategies for the genome assembly and annotation, which were applied by RYW. RYW and IvdH performed comparative analyses of Brassicaceae genomes. RH was involved in genome annotation. MES helped interpreting the results of the comparative analyses and was involved in drafting the manuscript. JH was involved in the design and interpretation of experiments that measured photosynthesis rates. FG, RYW, RB, DDR, MGMA, and SS were majorly involved in overall experimental design and preparing the manuscript. All authors read and approved the final manuscript.

## Acknowledgements

We gratefully acknowledge Pádraic Flood for collecting the *H. incana* “Nijmegen” accession and Graham Taylor for propagating it to perform the first gas exchange measurements. We thank Carlos Alonso Blanco for collecting the *H. incana* “Burgos” accession, Niccolò Bassetti and Nina Fatouros for collecting the *B. nigra* “DG2” accession. We thank the Unifarm staff at the Nergena greenhouse and Radix Klima climate chambers unit for taking excellent care of our plants. We thank Steven Driever for providing valuable insights on the modelling of light-response curves, José van de Belt for her help on chromosome counting, and the Applied Bioinformatics group of Wageningen Plant Research for performing RNA sequencing.

