## Additional File 1 for "The genome sequence of *Hirschfeldia incana*, a species with high photosynthetic light-use efficiency"

### Supplementary Methods

#### Pre-processing of reads

##### PacBio

Unless specified otherwise, we used the PacBio data in its native BAM format. For downstream applications which required FASTQ format, we converted the data from BAM to FASTQ using bam2fastq (version 1.3.0) (`https://github.com/PacificBiosciences/bam2fastx`).

### 10X

Unless specified otherwise, we used the 10X data in its native FASTQ format, which includes barcodes. The barcodes were removed using Long Ranger (version 2.2.2) (`https://support.10xgenomics.com/genome-exome/software/downloads/latest`) (longranger basic command), if downstream applications required it.

##### Illumina

We trimmed Illumina data using Cutadapt (version 1.18) [28]. Sequences were clipped from the 5' and/or the 3' ends of reads if they matched for at least 90% of their total length to one of the adapter sequences provided in the NEBNext Multiplex Oligos for Illumina (Index Primers Set 1). In addition, we trimmed bases which had a phred quality score of 30 or lower. Reads that were shorter than 100 bp after trimming were discarded. We removed duplicate Illumina read pairs using clumpify.sh of the BBTools suite (version 37.66) (`https://jgi.doe.gov/data-and-tools/bbtools`), allowing a maximum of three substitutions between putative duplicate pairs. The resulting deduplicated dataset was used for all downstream applications involving Illumina data.

#### Identifying nuclear, organellar, and contaminant reads

##### Identifying the origin of contigs in an initial assembly

We generated an initial genomic assembly of the PacBio data using Canu (version 1.8) [19] with the parameters genomeSize=325m and correctedErrorRate=0.040. As the PacBio data was generated from total-cellular DNA, the assembly contained contigs of mixed origin: nuclear, mitochondrial, chloroplast, or contaminant. To identify the origin of each contig, we aligned the assembly against the NCBI nucleotide (nt) database (obtained from `ftp://ftp.ncbi.nih.gov/blast/db` on 10-09-2019) using the command line version of blastn (version 2.6.0) [3], retaining the best hit of each contig. In addition, we investigated the coverage of each contigs by mapping PacBio reads against the assembly using pbmm2 (`https://github.com/PacificBiosciences/pbmm2`), a C++ wrapper of minimap2 [24] that can process PacBio reads in their native BAM format. Most contigs had hits against nuclear, mitochondrial, and chloroplast sequences of other plant genomes, primarily those of the Brassicaceae family. A small set of contigs had hits with genomes of contaminants such as mite and mite predator species, which act as pests or biological control, respectively. Consistent with known differences between relative copy numbers of nuclear and organellar DNA in plant species, contigs with nuclear hits generally had lower coverage (around 100x) than those with mitochondrial (900-1,100x) and chloroplast (at least 10,000x) hits. As such large differences in coverage can obfuscate the assembly process, we used both BLAST hits and coverage information to partition PacBio reads into different bins according to their origin.

##### Extracting chloroplast reads

We considered contigs to represent chloroplast sequence if they had excessive coverage of PacBio reads (at least 10,000x) and a BLAST hit to a plant chloroplast sequence in the nt database. We extracted alignments of reads to such contigs with samtools view (version 1.9) [25] and samtools collate, and recovered the original read sequences using bedtools bedtobam (version 2.27.1) [30]. To reduce coverage before proceeding with assembly, reads were subsampled to an estimated 200-300x coverage of the chloroplast genome with seqtk sample (version 1.2-r102-dirty) (`https://github.com/lh3/seqtk`).

##### Extracting mitochondrial reads

We considered contigs to represent mitochondrial DNA if they had 900-1,100x coverage of PacBio reads and a BLAST hit to a plant mitochondrial sequence in the nt database. The contigs that fulfilled these criteria had a combined length of 1 Mb, corresponding to about four copies of the mitochondrial genome. Therefore, we estimated that the reads that aligned to these contigs corresponded to about 4,000x coverage of the complete mitochondrial genome. We extracted these alignments, recovered the original read sequences, and subsampled reads to an estimated 200-300x coverage of the mitochondrial genome, using the same commands as described in *Extracting chloroplast reads.*

#### Assembling the chloroplast genome

##### Assembly

We assembled the chloroplast reads using Flye (version 2.5-release) [18] with the parameters –genome-size 200000 and –plasmid, yielding one single contiguous sequence. The assembly was polished by running one round of Arrow (version 1.0.0) (`https://github.com/PacificBiosciences/gcpp`), using alignments produced by pbmm2 of all raw PacBio reads to the assembly as input. The alignments were subsampled to 200-300x coverage with Picard DownsampleSam (version 2.9.2-SNAPSHOT) (`https://github.com/broadinstitute/picard`) to reduce running time.

##### Error correction

We performed detailed manual inspection to identify and correct errors in the chloroplast assembly. To check for correct circularization, we exchanged the positions of both halves of the contig and aligned the Illumina reads to it with bwa mem (version 0.7.10-r789) [23]. We subsampled the alignments to about 200x coverage with Picard DownsampleSam and called indels using Freebayes (version v1.3.1-dirty) [8] (parameters -p 1 -F 0.5 -m 50 –min-coverage 10 –read-mismatch-limit 5 –read-snp-limit 5 –limit-coverage 100) and GRIDSS (version 1.8.1-gridss) [4] (default parameters). This analysis revealed one 31 bp insertion in the middle of the sequence (the circularisation site of the original, unmodified contig) and two homozygous 1 bp indels outside of it, which were all corrected using bcftools consensus (version 1.9) [22].

##### Correcting orientation

It is conventional to report the sequence of plant chloroplasts in the following order: the long single copy (LSC) region, the first copy of the inverted repeat (IR), the short single copy (SSC) region, and finally the second copy of the inverted repeat region. We identified these regions in the assembly using the CPGAVAS2 web server [32] (accessed on September 20, 2019) and reordered the sequence of the contig accordingly to follow conventions. To check whether all regions were correctly oriented, we aligned our assembly to the chloroplast sequences of *A. thaliana*, *B. rapa*, and *B. napus* using minimap2 with parameter -ax asm20, revealing that the LSC of the chloroplast assembly of *H. incana* had been inverted relative to the LSC of the other three chloroplast assemblies. We manually inverted the LSC of the H. incana sequence to correct this error. We performed a final round of polishing with Arrow and FreeBayes/BCFtools (indels only) and repeated the check for correct circularisation, which resulted in the manual insertion of a single missing A at the end of the sequence.

##### Validation using an Illumina-based assembly

We generated an Illumina-based assembly of the chloroplast as an additional means of validation. Raw Illumina reads were assembled using IOGA [1], yielding one contig containing the LSC and IR, and one contig containing the SSC. These contigs were concatenated and the reverse complement of the IR was added at the end to create a complete chloroplast sequence. The sequence was polished by aligning Illumina reads to it with Bowtie2 [21] and using the resulting alignments as input to Pilon [38] and GapFiller [2] with default parameters. Any remaining stretches of Ns were filled by aligning the assembly to the *A. thaliana* chloroplast sequence with blastn and manually replacing the Ns by the *A. thaliana* sequence that aligned to them. Both halves of the assembly were exchanged to check whether the sequence properly circularized, resulting in the removal of 27 bp of sequence that overlapped between both ends. The final Illumina-based assembly was identical to the one based on PacBio reads, suggesting that both correctly represent the chloroplast sequence of *H. incana.*

##### Annotation

The chloroplast assembly was annotated using the cpGAVAS2 workflow [32] with default parameters and the 43 RNA-seq curated plastomes as reference set.

#### Assembling the mitochondrial genome

##### Assembly

We assembled mitochondrial reads using Canu with parameters genomeSize=250k and corrected-ErrorRate=0.040, yielding five contigs. Inspecting the assembly graph produced by Canu with Bandage (version 0.8.1) [39] revealed that two contigs formed a circular sequence, while the other three contigs corresponded to "dead-ends". The presence of multiple possible assembly paths is consistent with previous studies that have shown that plant mitochondrial DNA typically consists of a "master circle" confirmation containing the full mitochondrial sequence and multiple alternative subgenomic circular and linear conformations generated by repeat-induced recombination [34]. As it is conventional to report the "master circle" sequence [35], we mapped all the reads that were used as input to the five contigs with minimap2 (parameter -ax map-pb) and only retained those that aligned to the two contigs forming the circular sequence. Running Canu (parameters: genomeSize=250k, correctedErrorRate=0.040) on these reads produced one single contig representing the "master circle" conformation.

##### Error correction

We ran Circlator (–b2r_discard_unmapped) (version 1.5.5-docker2) [15] to correct any errors regarding circularization, using all mitochondrion-derived reads as input. We used the same set of reads to call structural variants with NGMLR (version 0.2.7) and Sniffles (version 1.0.9) [31] (parameters: -s 20). No structural variants were reported by this analysis, suggesting that the assembly was free of large misassemblies. We polished the assembly using two rounds of Arrow, aligning raw PacBio reads with pbmm2 and subsampling alignments to about 200-300x coverage with Picard DownsampleSam at each round. We manually assessed whether the assembly properly circularized by exchanging both halves of the sequence and mapping Illumina reads to it using bwa mem with default parameters. The chloroplast assembly of *H. incana* was included in the alignment procedure to prevent cross-mapping of chloroplast-derived reads. We used the alignments to call SNPs and indels with FreeBayes, resulting in a small set of small variants. However, all of them were located in regions which had relatively higher coverage than the rest of the genome. The nuclear genome of plants is known to contain inserted DNA that is highly similar to mitochondrial sequence and we suspect that the cross-mapping of such reads is what causes this rise in coverage. Thus, we consider any variants called in these regions to be incorrect. This result, combined with the absence of large structural variation based on mitochondrion-derived reads, suggests that our mitochondrial assembly is both accurate and complete.

#### Assembling the nuclear genome

##### Extracting nuclear reads

We attempted to remove non-nuclear reads from the PacBio data by mapping it with pbmm2 against the chloroplast and mitochondrial assembly (both duplicated to catch reads that span both ends), the reference genomes of any contaminants to which the contigs of the initial assembly had a BLAST hit, and contigs that had a BLAST hit with a contaminant. Unmapped reads in the output file were retained using the parameter –unmapped. A custom Python script based on the pysam library (`https://github.com/pysam-developers/pysam`) was used to extract all putative nuclear reads from the alignments. We considered reads to be of nuclear origin if they fulfilled any of the following conditions: the read was unmapped, the read had an alignment which was clipped by more than 5% of its length, or the read had an alignment which consisted for over 15% of its length out of mismatches. This latter threshold was chosen based on a histogram of the mismatch rate of all aligned reads, which showed a peak at about 10% (Figure S2), consistent with the known sequencing error rate of the PacBio Sequel platform [26]. Given that the organelles were sequenced at much higher depths than the nuclear genome, we considered reads which had such a mismatch rate to be derived from organelles and any reads that had a mismatch rate higher than this peak (over 15%) to be nuclear-derived.

##### Assembly

We assembled nuclear reads using Canu (parameters: genomeSize=325m, correctedErrorRate=0.040). To remove misassemblies, we identified and broke putative misassemblies of the Canu assembly with Tigmint (version 1.1.2) [16], which uses alignments of 10X reads as input. Alignments were produced using bwa mem with default parameters on 10X data processed by Long Ranger. Besides the assembly produced by Canu, the organellar assemblies and reference genomes of possible contaminants were included in the alignment procedure to prevent organellar and contaminant reads from introducing false misassemblies. Alignments were sorted by barcode using samtools sort (parameter -tBX).

##### Quality control

We assessed the quality of the assembly through several measures. We determined the contiguity of the assembly using QUAST (version 5.0.2) [11]. To estimate the completeness of the genic space, we aligned RNA-seq reads to the assembly with HISAT2 (version 2.1.0) [17]. We executed BUSCO (version 3.0.2) [33] to determine how well the assembly covered the embryophyta_odb9 database of conserved plant orthologs. To identify erroneously assembled contigs, we mapped the putative nuclear-derived PacBio reads to it with minimap2 (-ax map-pb) and computed the read depth of each contig with mosdepth (version 0.2.6) [29]. Finally, we identified possible contaminant contigs by aligning the Canu assembly against the nt database with blastn (-max_target_seqs 1 -max_hsps 1 -evalue 1e-25).

##### Filtering contigs

The results of the quality control step were used to design a set of filters for obtaining contigs that are expected to be reasonably accurate. We kept contigs that were covered for at least 90% of their length by at least one PacBio read, if they had no BLAST hit with a non-plant derived sequence, and fulfilled any of the following conditions: having a median read depth of at least 30X, containing a BUSCO, or having RNA-seq coverage.

##### Removing duplicate haplotigs and scaffolding

We purged alternative haplotigs from the assembly using purge_dups (version 1.0.0) [10] with default parameters. The purged assembly was scaffolded with ARKS (version 1.0.4) [5] (parameters c=8 m=8-10000), a tool which infers linkage between contigs using 10X data.

##### Gap-filling and polishing

After performing scaffolding, we filled gaps and polished the assembly with Arrow, using alignments of raw PacBio reads as input. The organellar assemblies and earlier identified contaminant and filtered contigs were included in the reference file during alignment to prevent reads derived from these sequences from obfuscating the polishing step. To remove any remaining single nucleotide level errors, we aligned 10X reads to the polished assembly using bwa mem, again including organellar, contaminant and filtered sequences in the reference file to prevent cross-mapping. We called variants in the assembly using FreeBayes (-p 2 -F 0.5 -m 50 –min-coverage 10 –read-mismatch-limit 5 –read-snp-limit 5 –limit-coverage 100) and corrected all indels that were well supported using bcftools consensus ('QUAL>1 && TYPE="indel" && (GT="AA" || GT="Aa")').

##### Filtering scaffolds

We used the same quality control steps as described in the *Quality control* section to assess the accuracy of all polished scaffolds. As an additional means of validation, we mapped 10X reads and Illumina to the polished scaffolds, organellar, contaminant and filtered sequences using respectively Long Ranger (longranger align) and bwa mem with default parameters. We designed another set of filters to only retain scaffolds which we deemed to be highly accurate. Scaffolds were kept if they contained a BUSCO or at least 10000 bp of RNA-seq coverage. Of the set that remained, we excluded scaffolds of which fewer than 90% of the length had a coverage of at least 6x based on Illumina reads, no BLAST hit with a plant sequence of the nt database, or fewer than 30x and 25x median coverage of PacBio on Illumina reads, respectively.

##### Removing sequences masked during duplicate haplotig removal

The assembly contained regions of 23 Ns, which were introduced by purge_dups while removing duplicated haplotig sequences. Although we were not able to ascertain that all sequences connected by these regions should be linked, there was evidence that some of the links were correct as breaking the scaffolds at these regions resulted in a loss of complete BUSCOs. Therefore, we decided to keep most of these regions in the assembly, only removing those that were present within 5 kb of scaffold ends, along with the associated sequences present at those ends.

##### Removing organellar sequences

To check for organellar sequences, we aligned the nuclear assembly against the mitochondrial and chloroplast assembly using blastn (-max_target_seqs 10 -max_hsps 10 -evalue 700). This revealed four nuclear regions which had near 100% identity with regions of the mitochondrial assembly and had significantly higher coverage of Illumina reads compared to the genomic average. We considered these regions to be mitochondrial and removed them from the nuclear assembly.

#### Genome annotation

##### Repeat annotation

RepeatModeler2 [7] (run with the -LTRstruct parameter) was used to generate a de novo library of repetitive elements for the *H. incana* nuclear genome, after which RepeatMasker [36] was used to soft-mask those elements. Both tools were called through TETools v1.1 (`https://github.com/Dfam-consortium/TETools`).

##### Gene annotation

We annotated putative protein-coding genes in the nuclear assembly using EvidenceModeler (commit 5d9425a, `https://github.com/EVidenceModeler/EVidenceModeler`) [13], which performs a weighted combination of de novo gene predictions, protein alignments, and transcript assemblies.

The input for EvidenceModeler was generated as follows. We aligned RNA-seq reads to the nuclear and organellar assemblies of *H. incana*, as well as against filtered contigs and contigs of possible contaminants. Reads that mapped against the nuclear assembly were extracted and used along with the masked nuclear genomic assembly as input to BRAKER (version 2.1.5) [14], which internally calls Genemark-ET (in RNA-seq guided mode) [27] and Augustus [37] to predict de novo gene models. We assembled transcripts out of the RNA-seq read alignments using Stringtie2 (version 2.1.1) [20] and detected ORFs in these transcripts using transdecoder (version 5.5.0) (`github.com/TransDecoder/TransDecoder`), both with default parameters. We obtained protein sets of *A. thaliana* (`www.arabidopsis.org/download_files/Proteins/Araport11_protein_lists/Araport11_genes.201606.pep.fasta.gz`), *B. rapa* (`brassicadb.org/brad/datasets/pub/Genomes/Brassica_rapa/V3.0/Brapa_genome_v3.0_pep.fasta.gz`), *B. nigra* (longest isoforms of the Ni100 annotation, translated with gffread), and the Viridiplantae clade (SwissProt set only: `www.uniprot.org/uniprot/?query=reviewed:yes%20taxonomy:33090`). All four sets of proteins were aligned to the *H. incana* nuclear genome using GenomeThreader (version 1.7.3) [9] (parameters: -species arabidopsis -gff3out -skipalignmentout).

The output of the two gene predictors, the four sets of protein alignments, and the ORFs predicted by transdecoder were given as input to EvidenceModeler to generate consensus gene models. These models were then processed with PASA (version 2.4.1) [12] to add UTRs and define alternative transcripts for each gene. PASA was ran by first aligning the transcripts assembled by Stringtie2 to the *H. incana* nuclear genome with GMAP and generating maximum alignment assemblies out of the alignments, followed by two rounds of updates to the EvidenceModeler consensus models. The final output of PASA was used for further filtering steps.

##### Filtering gene annotation

We filtered transcripts based on homology with proteins with close relatives and support from RNA-seq reads. We extracted the longest isoforms of the annotated genes in *A. thaliana*, *B. rapa* (v3.0 assembly), and *B. nigra* (Ni100 v2 assembly) using AGAT (version 0.2.3) (`github.com/NBISweden/AGAT`), converted them into protein sequences using gffread (v0.11.8) (`https://github.com/gpertea/gffread`), and clustered them with the protein products of *H. incana* into orthogroups using OrthoFinder (version 2.3.11) [6]. The *A. thaliana* annotation used was a combination of the TAIR10 genomic annotation of Ensembl Plants (release version 40) and the annotation of an improved A. thaliana mitochondrial assembly (BK010421.1).

We extracted the exons of the *H. incana* annotation in BED format and used mosdepth (version 0.2.9) (ran with the parameter -T 1) to determine which exons were supported by at least one RNA-seq read. We extracted intron-spanning reads of the RNA-seq alignments and introns from the *H. incana* annotation using custom Python scripts and intersected them with bedtools intersect to compute the support for each intron.

Transcripts were then filtered using the following the criteria:

- All introns supported by at least one RNA-seq read, at least 85% reciprocal overlap with a transcript assembly produced by Stringtie2
- Overlap with a protein alignment of *A. thaliana*; *B. rapa*, or the Viridiplantae SwissProt dataset for at least 85% of the length of the protein alignment and at least 25% of the length

of the transcript (the length threshold for the transcript is less strict, because a protein alignment lacks UTRs);

- Has a homolog with an *A. thaliana* gene involved in photosynthesis.

Transcripts which fulfilled at least one of these three criteria were put in the "accepted" set and those that did not were put in the "rejected" set. As a final filtering step, we removed transcripts of the "accepted" set which overlapped for over 85% of their length with repetitive elements identified by RepeatMasker, as these are likely to be involved with transposable elements.

*A. thaliana* genes were considered to be involved in photosynthesis, if they fulfilled one of the following conditions:

- Assigned one of the following GO terms: "photosynthesis", "electron transporter, transferring electrons within the cyclic electron transport pathway of photosynthesis activity", or "electron transporter, transferring electrons within the noncyclic electron transport pathway of photosynthesis activity";
- Included in the KEGG pathways ath00195 (Photosynthesis), ath00710 (Carbon fixation in photosynthetic organisms), and ath00196 (Photosynthesis - Antenna Proteins);
- Protein products have been assigned the keyword "Photosynthesis" in the Swiss-Prot database.

##### Assessing completeness through protein alignments of *B. rapa*

The longest isoforms of *B. rapa* genes were obtained using AGAT and translated to proteins with gffread. Translated proteins were aligned to the *H. incana* assembly using GenomeThreader (parameters: -species arabidopsis -gff3out -skipalignmentout), that reports matches if proteins align to the assembly for at least 50% of their length. 31002 (67%) of the *B. rapa* proteins could be aligned to the *H. incana* assembly in this manner, resulting in a total of 37283 alignments.

#### Supplementary Figures

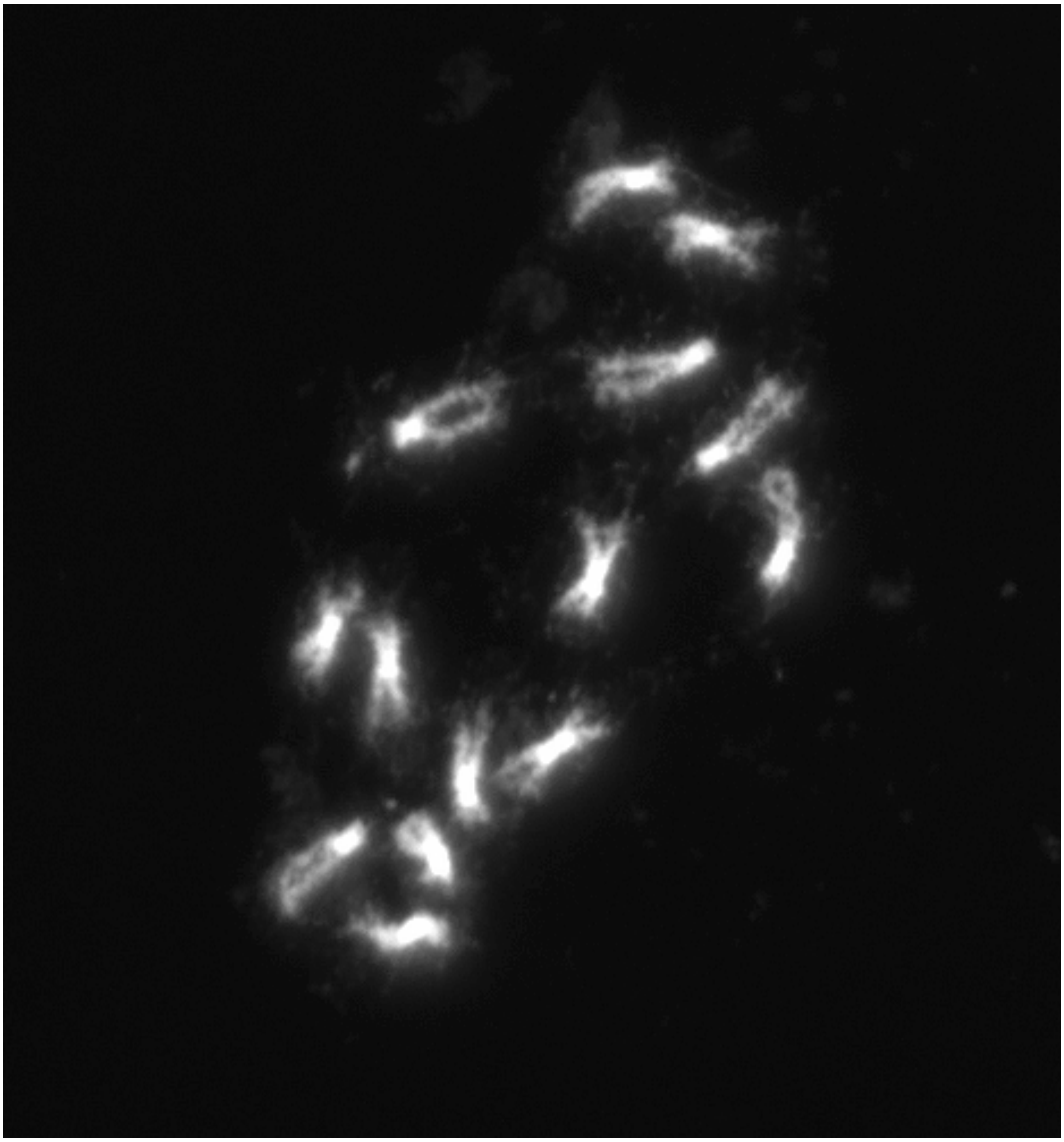

Figure S1: **Root tip smears of *H. incana*.** An example of DAPI-stained chromosomes from one metaphase mitotic cell. Fourteen chromosomes can be observed, meaning that *H. incana* has a haploid chromosome number of $n = 7$.

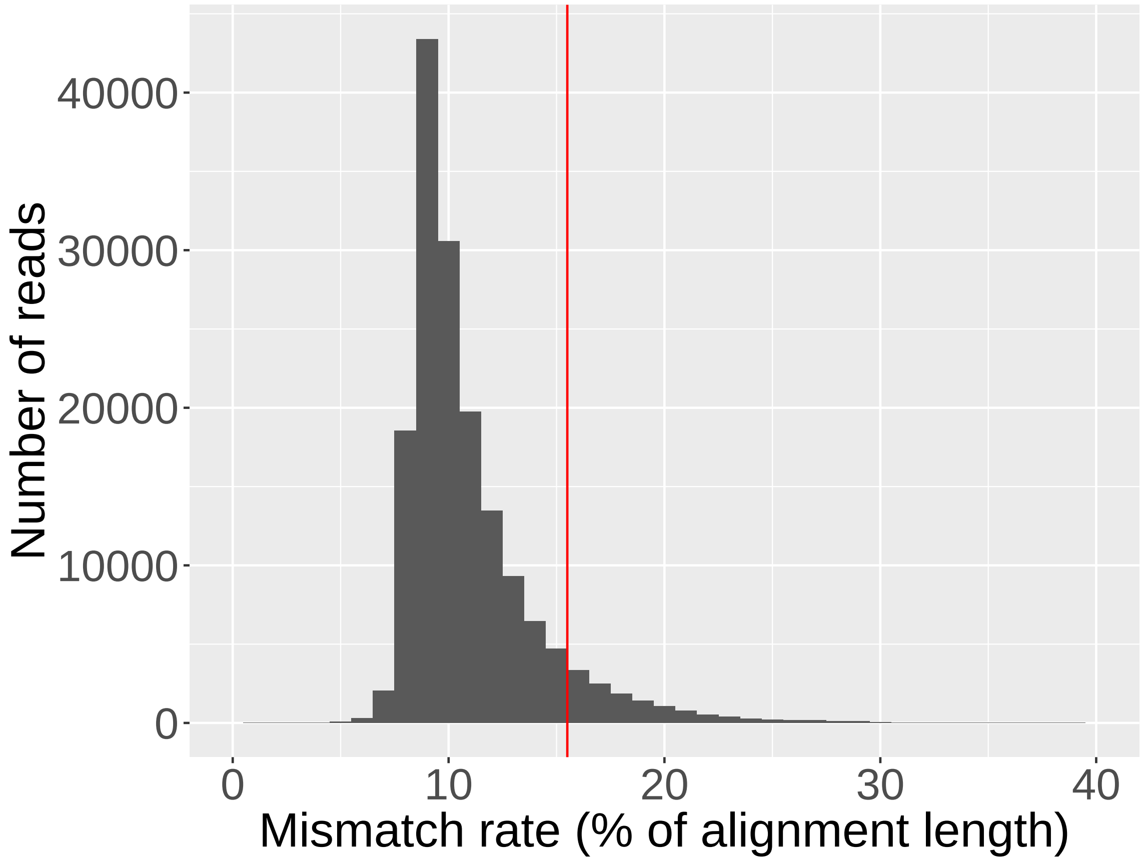


Figure S2: **Histogram of mismatch rates of PacBio reads aligned against the chloroplast and mitochondrial assembly of *H. incana*.** The cut-off of 15%, used to distinguish organelle-derived reads from reads originating from nuclear regions that contain organellar inserts, is depicted by the red vertical line.
